# A VSV-vector vaccine simultaneously targeting H5N1 hemagglutinin (HA) and matrix protein 2 (M2) induces robust neutralizing and ADCC antibody responses and provides full protection against lethal H5N1 infection in a mouse model

**DOI:** 10.64898/2025.12.26.696590

**Authors:** Zhujun Ao, Robert Vendramelli, Maryann Buyu, Thang Truong, Dayu Liu, Nick Gao, Stephen Amos, Peter Lawrynuik, Maurice Nyamekye, Keith R. Fowke, Darwyn Kobasa, Xiaojian Yao

## Abstract

Human (avian) influenza A viruses, especially highly pathogenic avian influenza (HPAI) viruses, pose a significant public health threat, and a multivalent vaccine is the primary prophylactic measure to control these viruses. To establish such a vaccine, we generated two multivalent vesicular stomatitis virus (VSV)-based vaccine candidates (V-EtM2e/H5_05_ and V-EtM2e/H5_22_) and characterized their ability to induce protective immune responses. Our results revealed that vaccine immunization in mice induced high humoral immune responses against both the HPAI hemagglutinin (HA) protein and the ectodomain of M2 (M2e) protein. Intriguingly, vaccine-immunized mouse sera exhibited highly efficient neutralizing activity against the corresponding H5 pseudovirus and mediated potent and broad antibody-dependent cellular cytotoxicity (ADCC) activity against M2e derived from human and avian influenza H5, H1, H3, and H7 viruses. Furthermore, both intranasal and intramuscular immunization provided efficient protection against HPAI H5N1 virus challenge in mice, with a 100% survival rate and a nondetectable viral load in several tissues. Notably, noninvasive mucosal (IN) delivery of V-EtM2e/H5_22_ achieved protection equal to that of IM delivery at a 100-fold lower immunizing dose. These findings provide strong evidence for the effectiveness of a multivalent VSV-based vaccine against human (avian) influenza A viruses.

## Introduction

Influenza is a highly contagious airborne disease that primarily affects the respiratory tract, causing recurrent seasonal epidemics and, at times, catastrophic pandemics. Among influenza A viruses, avian influenza viruses (AIVs) are particularly concerning because of their extensive circulation in domestic and wild bird populations and their ability, under certain conditions, to cross the species barrier into mammals, including humans^1,2^. Avian influenza viruses are categorized as either low pathogenic avian influenza (LPAI) or highly pathogenic avian influenza (HPAI) viruses. LPAI viruses often cause mild or asymptomatic infections in poultry^2^, whereas HPAI viruses induce severe disease with high mortality^2,3^. H5N1 is the most widely studied HPAI subtype. First identified in Guangdong, China, in 1996 as A/goose/Guangdong/1/96^4^, H5N1 has rapidly spread across Asia, Europe, the Middle East, Africa, and even North America^5,6^. Large outbreaks in migratory and wild birds have established long-term viral reservoirs, increasing the likelihood of extension to agricultural systems and human populations. In 2024, H5N1 caused a multistate outbreak in poultry and dairy cows in the US, resulting in human infections primarily affecting those with occupational exposure to infected animals ^7^. Although sustained human-to-human transmission of this virus has not been documented, viral evolution through antigenic drift or reassortment with seasonal influenza A viruses could confer this property, making H5N1 a leading candidate for causing a pandemic. Therefore, developing vaccines for HPAI viruses is a high public health priority to combat a potential epidemic or HPAI pandemic.

Despite this urgency, human HPAI vaccine development remains in its early stages. Current influenza vaccines, which largely rely on hemagglutinin (HA) ^8^ or neuraminidase (NA) ^9^ antigens, provide strain-specific neutralizing protection and are not universal ^10–12^, leaving individuals vulnerable to new or divergent AIV subtypes. Furthermore, the traditional egg-based manufacturing process is unsuitable for HPAI vaccine production because of the virus’s high pathogenicity in eggs, which adds barriers to scale and costs ^13^. Thus, two central challenges in HPAI vaccine design are improving multivalent efficacy across diverse viral strains and reducing production costs to enable widespread access. Addressing these challenges requires novel approaches that target conserved viral components, with the ultimate goal of developing universal influenza vaccines capable of eliciting broad, durable, and affordable protection against a wide spectrum^14,15^ of influenza A viruses.

In addition to antigen design, the choice of vector plays a pivotal role in vaccine efficacy. The recombinant vesicular stomatitis virus (rVSV) vector has several advantages, including highly efficient delivery of antigens to host cells and the induction of strong humoral and cellular immune responses. Its broad host range, capacity for robust replication, and ability to be engineered for safety make rVSV an attractive platform for vaccine development. Importantly, rVSV-based vaccines have demonstrated the ability to induce durable protective immunity and can be adapted for both intramuscular and mucosal delivery, thereby supporting their versatility for controlling emerging viral threats.

Our group previously developed a recombinant vesicular stomatitis virus (rVSV)-based universal influenza vaccine candidate incorporating four tandem repeats of the influenza M2 ectodomain (tetra-M2e, tM2e) fused to the dendritic cell-targeting domain of Ebola virus glycoprotein (EΔM) (rVSV-EboΔ-tM2e) ^16^. This design aimed to leverage the high sequence conservation of M2e to induce antibody-mediated cellular events such as antibody-dependent cellular cytotoxicity (ADCC) (ref), antibody-dependent cellular phagocytosis (ADCP) (ref), and complement-dependent cytotoxicity (CDC) (ref), thereby broadening protection against diverse influenza A subtypes. Our results demonstrated that the rVSV-EΔM-tM2e vaccine induced rapid and potent M2 antibody production and ADCC activity and protected mice from H1N1 and H3N2 influenza challenges. In this study, we sought to address these limitations by developing a VSV-based multivalent vaccine designed to target two distinct influenza membrane antigens, the HA from a recently isolated HPAI H5N1 strain (Influenza A/Red-Tailed Hawk/ON/FAV-0473-4-2022), and four copies of the ectodomain of influenza M2 (human/bird/swine) that fused with an Ebola virus glycoprotein dendritic cell-targeting domain (EΔM). Influenza HA is the primary target of neutralizing antibodies, yet its high variability limits cross-protection. By combining HA with the highly conserved M2e antigen, this vaccine aims to elicit both potent neutralizing antibody responses against HA and antibodies that mediate ADCC against highly conserved regions of the influenza M2 ectodomain to provide broad protection against HPAI viruses. The use of the VSV vector and EΔM for dendritic cell targeting further enhances antigen presentation and immune priming. Collectively, this approach integrates antigenic breadth with potent immune activation, offering a promising strategy toward a more universal influenza vaccine.

## Materials and Methods

### Plasmid constructs

To generate the rVSVΔG vector coexpressing both the codon-optimized genes encoding avian influenza H5N1 hemagglutinin (HA) and EΔM-tM2e (EtM2e) ^16,17^, H5_05_ (Influenza A/Hanoi/30408/2005 (H5N1), NCBI accession No. AB239125.1) ^18^ or H5_22_ (Influenza A/Red-Tailed Hawk/ON/FAV-0473-4-2022 (H5N1), EPI_ISL_17394087) ^19^ were cloned and inserted into a VSVΔG vector at the position of the VSV-G gene using a previously described cloning strategy ^20^, and the resulting vaccine candidates were named V-EtM2e/H5_05_ and V-EtM2e/H5_22_.

The pCAGGS-DNAs encoding HA(H5_05_), neuraminidase (NA) (NCBI accession no. AB239126.1), and M2_05_ (A/Hanoi/30408/2005) were previously described ^21^. cDNA encoding M2e from H7N9 (human and avian) was generated by two-step PCR and subsequently cloned and inserted into the pCAGGS-M2 expressor. The codon-optimized gene encoding HA(H5_22_) was synthesized by GenScript Biotech, Inc., and cloned and inserted into the pCAGGS vector. The protease cleavage site of H5_22_ was modified by a two-step PCR technique to delete five or four basic amino acids (RRRKK for H5_05;_ KRRK for H5_22_) and add a threonine residue proximal to the cleavage site of the protein (Fig. 1B), as previously described ^21^. The lentiviral vector encoding the luciferase gene was previously described ^22^.

**Figure 1.**
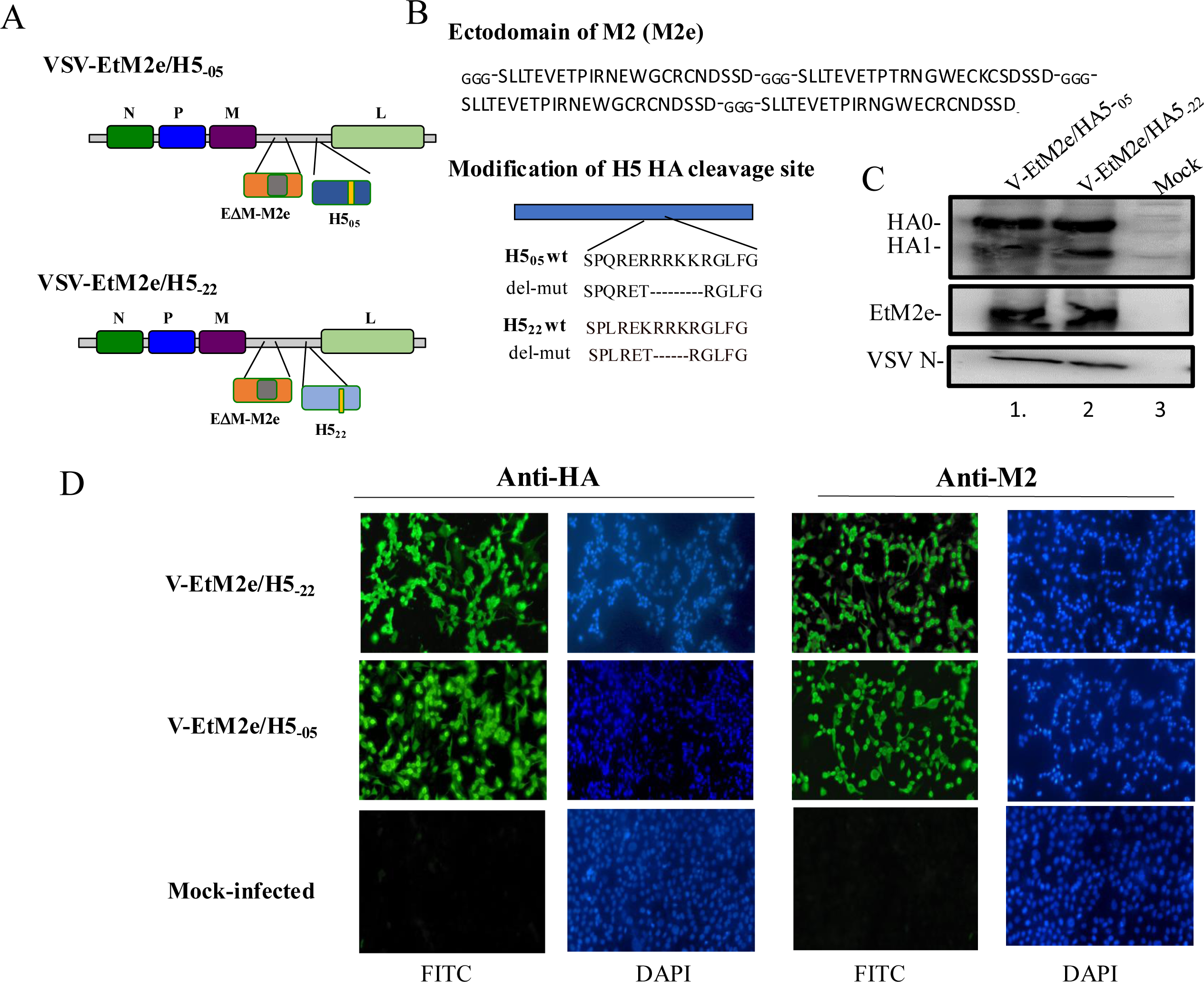
Construction and expression of V-EtM2e/H5_05_ and V-EtM2e/H5_22_ multivalent vaccines. A) Schematic diagram of the EΔM-tM2e, H5_05,_ or H5_22_ immunogens present in the VSV vaccines. EΔM-tM2e, four copies of influenza virus M2 ectodomain (24 aa) polypeptide (tM2e) inserted into the EΔM. B) Four copies of the conserved influenza M2 ectodomain (24 aa) polypeptide (tM2e, comprised of two copies from human flu strains, one from swine flu strain and one from avian flu strain (top panel), In H5_05_ and H5_22_ open reading frames, a deletion was made to convert the furin-dependent cleavage site to a trypsin-dependent activation site (low panel). C) VeroE6 cells infected with V-EtM2e/H5_05_ or V-EtM2e/H5_22_, were lysed and processed with SDS-PAGE followed by WB with a rabbit anti-HA antibody (top panel), a mouse antibody against influenza M2e (middle panel) or mouse anti-VSV nucleocapsid (N) (low panel). D) Representative immunofluorescence images of Vero E6 cells infected with V-EtM2e/H5_22_ (top panel), V-EtM2e/H5_05_ (middle panel) or mock-infected (low panel). Infected cells were incubated with anti-HA antibody or anti-M2e antibody, followed by corresponding FITC-conjugated secondary antibodies. Cells were viewed under a computerized Axiovert 200 inverted fluorescence microscopy.

### Ethics statement

The animal experiments described were carried out according to protocols approved by the Central Animal Care Facility, University of Manitoba (Protocol Approval No. B2024-030/1), following the guidelines provided by the Canadian Council on Animal Care. All animals were acclimated for at least one week before experimental manipulation and given ad libitum access to food and water in a specific pathogen-free animal facility.

All animal challenge experiments carried out at the National Microbiology Laboratory (NML) of the Public Health Agency of Canada were approved by the Animal Care Committee at the Canadian Science Center for Human and Animal Health per the guidelines provided by the Canadian Council on Animal Care under animal use document H-23-004. The mice were monitored by registered animal health technicians throughout the experiment and given ad libitum access to food and water. All infection work with the highly pathogenic avian influenza (HPAI) A(H5N1) was performed in the containment level 4 (CL-4) laboratory at the NML. The animals were acclimatized for 5–7 days before the beginning of all the experiments. All the animals were monitored and weighed daily throughout all the experiments and given ad libitum access to food and water. All manipulations, such as blood collection and infection, were performed under isoflurane anesthesia.

### Cells, plasmids, antibodies, recombinant proteins, and viruses

The HEK293T, VeroE6, and BHK-T7 cell lines were cultured in DMEM. The ADCC bioassay effector Jurkat cell line (BPS Bioscience, Cat# 78733) was cultured in RPMI-1640 medium (ATCC modification). Madin-Darby Canine Kidney (MDCK) cells obtained from the American Type Culture Collection (ATCC) were cultured in minimal essential medium (MEM) (Gibco) supplemented with 5% bovine growth serum (HyClone) and 1% penicillin and streptomycin (P/S; 100 units/mL and 100 µg/mL, respectively; Gibco cat# 15140-122) for maintenance. The antibodies used in this study included a rabbit polyclonal antibody against HA (Sino Biological, Cat# 11700-T54), a mouse monoclonal anti-influenza A virus M2 antibody (Santa Cruz Biotech, Cat# 14C2: sc-32238), and a mouse monoclonal anti-VSV-N antibody (EMD Millipore Corporation, Cat# MABF2348). The secondary antibodies used were anti-mouse IgG-HRP (GE Healthcare, Cat# NA931), goat anti-mouse-IgG (H+L)-HRP, goat anti-mouse-IgG2a-HRP, and goat anti-mouse-IgG1-HRP (Southern Biotech, Cat# 1037-05, 1081-05, or 1071-05). The recombinant proteins included influenza A H5N1 HA (A Cambodia/NPH230032/2023, Sino Biological, Cat# 409450-V08B), H5N1 HA (A/bald eagle/Florida/W22-134-OP/2022, BEI Resources, Cat# NR-59424), H5N2 HA (A/snow goose/Missouri/CC15-84A/2015, BEI Resources, Cat# NR-50651), H7N9 HA (A/Human/02285/2017, BEI Resources, Cat# NR-51195), H3N2 HA (A/Missouri/09/2014, BEI Resources, Cat# 40494-V08B) and H1N1 HA (A/Puerto Rico/8/34, Sino Biological, Cat# 11684-V08H). The cDNA encoding the EboGPΔM-tM2e fusion was previously described ^16^.

The replication-competent recombinant vesicular stomatitis virus (rVSV)-based influenza vaccines V-EtM2e/H5_05_ and V-EtM2e/H5_22_ were rescued in Vero E6 cells, propagated, and titrated as described previously ^20^. The influenza A H5N1 virus A/BC/PHL_2032/2024 (GISAID # EPI_ISL_19548836) (A/BC(H5N1)) used to infect mice in the challenge experiments at the NML, Winnipeg, was isolated from a human with nonfatal infection in Canada ^23^.

### Western blot and immunofluorescence assays

To detect the expression of different viral proteins, Vero E6 cells were infected with V-EtM2e/H5_05_ or V-EtM2e/H5_22_ for 48 hours, and the infected cells were lysed and subjected to 10% SDS‒PAGE, followed by immunoblotting with various antibodies, including anti-HA, anti-M2 or anti-VSV-N. The protein bands were visualized using an enhanced chemiluminescence kit (Perkin Elmer Life Sciences).

To detect the ability of the serum of vaccinated mice to M2 from different strains, 293T cells were transfected with plasmids encoding M2 from different strains for 48 hours. The cell lysates were then run on a 12% SDS‒PAGE gel, followed by immunoblotting with a V-EtM2e/H5_22-_immunized mouse serum pool.

For the indirect immunofluorescence assay, Vero E6 cells grown on glass coverslips (12 mm^2^) in 24-well plates were infected with V-EtM2e/H5_05_ or V-EtM2e/H5_22_ for 24 hours. After infection, the cells on the coverslip were fixed with 4% paraformaldehyde for 5 minutes and permeabilized with 0.2% Triton X-100 in PBS. The cells were then incubated with primary antibodies specific for M2e or HA, followed by the corresponding secondary FITC-conjugated antibodies. The cells were finally viewed under a computerized Axiovert 200 inverted fluorescence microscope.

### Mouse immunization and viral challenge

For the immunization experiments, female BALB/c mice aged 6 to 8 weeks (five per group) were immunized IM (1×10^7^ TCID50) with VSV the vaccine candidates (V-EtM2e/H5_05_ or V-EtM2e/H5_22_) and boosted on day 14. At 28 days post-immunization, the mice were sacrificed, and blood samples were collected for various analyses.

For virus challenge and sample collection, eighty BALB/c mice were purchased from Charles River Laboratories. Upon arrival, the mice were transported to a containment level 2 (CL-2) animal facility, where they were separated by sex with 5 in each cage and allowed to acclimate for 7 days prior to vaccination. The mice were assigned to one of two vaccination arms for intranasal (IN) or intramuscular (IM) delivery, with 40 mice in each arm. These vaccination arms were then divided in half again to develop vaccinated groups and control groups. The final groups included 20 mice of equal sex. During IN vaccinations, the mice were placed under isoflurane anesthesia, and the animals were primed with 1 × 10^5^ TCID_50_ in 50 µL of V-EtM2e/H5_22_ vaccine (IN vaccination group) or 1 × 10^5^ TCID_50_ in 50 µL of VSV-EBOVGP (IN control group), both of which were delivered via intranasal instillation. In the IM vaccination arm, mice were placed under isoflurane anesthesia and primed with a total of 1 × 10^7^ TCID_50_ of the V-EtM2e/H5_22_ vaccine delivered in two 50 µL injections into each hind leg (IN vaccination group), while the IM control group was primed in the same manner with 1 × 10^7^ TCID_50_ of VSV/EBOVGP. All the animals were boosted at 21 days after prime vaccination (dpv) using the same method. Serum was collected from 6 mice (3 males and 3 females) prior to priming and from all the mice at 20 dpv and 41 dpv.

Following serum collection at 41 dpv, the mice were transferred to the containment level 4 (CL4) facility at the National Microbiology Laboratory (Winnipeg, MB, Canada). Mice were challenged intranasally (IN) on day 42 under isoflurane anesthesia with 10× LD_50_ (10 PFU) of a highly pathogenic H5N1 strain (A/BC/PHL_2032/2024) in a total volume of 50 µL (25 µL per naris). The animals were weighed and monitored daily and euthanized if they were assigned a preapproved clinical score. Half of the animals were necropsied at 4 days post-infection (dpi), and serum, lung, heart, kidney, and spleen samples were collected in 2 mL cryovials. The tissue samples were weighed and flash frozen at −80 °C for further downstream analysis. All other mice were monitored until 18 dpi.

### Measurement of viral burden in the tissues

To measure the viral titers in the tissues that were collected at 4 dpi, a median tissue culture infectious dose (TCID_50_) assay was performed. MDCK cells were seeded into 96-well tissue culture plates as described previously ^20^ and brought into CL4 for the addition of tissue homogenate dilutions. Tissues previously collected from euthanized animals were thawed and homogenized with 5 mm sterile stainless steel beads using a Bead Ruptor Elite Tissue Homogenizer (Omni). The homogenates were clarified by centrifugation at 1500 × g for 10 minutes, after which tenfold dilutions of the tissue homogenates were made in viral growth media. Dilutions were added to MDCK cells in triplicate wells, and the cytopathic effect was assessed at 72 hours post-infection. TCID_50_ values per gram of tissue were calculated using the Reed and Muench method.

### Measurement of vaccine-induced HA- and M2e-binding antibody titers by ELISA

To determine the HA- and M2e-specific antibody titers in immunized mouse serum samples, ELISA was performed as previously described ^24^. Briefly, 96-well plates (NUNC MaxiSorp, Thermo Scientific) were coated with recombinant HA (H5N1, A/Cambodia/NPH230032/2023, Sino Biological, Cat# 40945-V08B) or M2e (human, GenScript, Cat# RP20206) (1 µg/mL) proteins in coupling buffer (pH 9.6, 50 mM sodium carbonate–bicarbonate) at 4 °C overnight. After blocking at 37 °C with blocking buffer (1% BSA (w/v) in 1× TBS) for 2 hours, the ELISA plates were washed and incubated with 2× serially diluted mouse serum samples at 37 °C for 1 hr. Goat anti-mouse-IgG(H+L)-HRP, goat anti-mouse-IgG2a, or goat anti-mouse-IgG1 (SouthernBiotech) was used to detect the antibodies binding to HA or M2e. After the samples were incubated with the substrate tetramethylbenzidine (TMB) solution (Mandel Scientific) and terminated with stop solution, the absorbance of each well at 450 nm (OD450) was measured. The endpoint titers of mouse serum samples were calculated using interpolation in GraphPad Prism 10.0, with a cutoff of 2.5 times the mean negative.

### HA pseudovirus preparation and Neutralization assay

H5N1 HA-pseudotyped Luciferase (luc)-expressing viral particles (H5-Luc-pseudovirus) were produced by co-transfecting 293T cells with avian influenza envelope expression plasmids (HA_05_ or HA_22,_ NA, or M2), the HIV packaging plasmid pCMVΔR8.2 and a lentiviral vector expressing Luciferase (pEF1-Luc/puro) ^22^. After 48 h of transfection, the supernatant containing H5-pseudoviruses will be filtered through a 0.45 μm filter and kept in −80°C.

The neutralization assay was performed based on influenza H5-pseudovirus and 293T cells according to the previous method ^21^, with some modifications. Briefly, the H5-pseudovirus were pre-activated by trypsin treatment (100 μg/ml) for 30min at 37°C before infection. Then the trypsin activated H5-pseudoviruses (about 10^4^ RLUs, 25 µL) were pre-incubated with 2× serially diluted inactivated mouse sera (starting from 1:20 dilution, 25 µL) in complete DMEM in a 96-well plate at 37 °C for 1 h. Then 293T cells (1.5×10^4^ cells/well, 50 µL) were added in each well and incubated at 37 °C. After 48-72 hrs of infection, cells were lysed in luciferase lysis buffer (Promega; 30 μL/well) and the luciferase relative light unit (RLU) in the lysates was measured using Polerstar optima microplate reader (BMG BioLabtech) according to the manufacturer’s instructions. The neutralizing titers or half-maximal inhibitory dilution (ID_50_) were defined as the reciprocal of the serum maximum dilution that reduced 50% in RLUs compared with no-serum controls. The ID_50_ was calculated by using sigmoid 4PL interpolation with GraphPad Prism 10.0.

### Antibody-dependent cellular cytotoxicity (ADCC) reporter assay

The ADCC reporter assay was performed by using ADCC Bioassay effector Jurkat cell line which expresses firefly luciferase under the control of NFAT (nuclear factor of activated T cells) response elements, and mouse FcγRIV (BPS Bioscience). The engagements among FC portion of antibodies (serum) with mFcγRIV on cell surface of the Jurkat effector cells and antigen-expressing target 293T cells activate the luciferase expression driven by NFAT in effector cells. Briefly, one day before the assay, 293T cells (6×10^5^/well, 6-well plate) were transfected with each plasmid expressing HA_05_, HA_22_ or M2 which carrying the M2e of influenza H5N1 or H7N9 using Lipofectamine 2000 (Invitrogen, USA) to produce 293T-HA or 293T-M2 target cells. The next day, the immunized mouse sera were 3× serially diluted in Assay medium 2A (BPS Bioscience#79621) (starting from 1:30, 50 µL/well) in a 96-well plate, then mixed with target cells (1×10^4^, 50 µL/well), incubated at 37 °C for 1 hour. The Jurkat effector cells (6×10^4^, 50 µL/well) were added and cultured for 5-6 hrs or overnight at 37 °C with 5% CO2. Finally, 150ul of one-step luciferase reagent (BPS Bioscience, Cat#60690) was added to each well and the ADCC activities were determined by measuring the Luciferase from activated Jurkat cells using a Polerstar optima microplate reader. The reading of the background well (only medium) was deducted. The no-serum controls were wells containing target cells and Jurkat cells. The fold of change (ADCC induction) was calculated as RLU (test − background)/RLU (no-serum control − background).

### Statistics

Statistical analysis of antibody levels was performed using an unpaired t test (with P ≥ 0.05 considered to indicate statistical significance) with GraphPad Prism 10.0 software. Statistical analysis of neutralizing antibodies and ADCC activity was performed using one-way ANOVA with multiple comparison tests followed by Tukey’s test in GraphPad Prism. Moreover, two-way ANOVA with multiple comparison tests was used to analyze mouse survival data, and the Mantel‒Cox test was used to analyze tissue TCID data.

## Results

### 1. Construction and characterization of a VSV vaccine expressing a modified H5N1 HA protein and four EboΔM-fused copies of influenza M2 ectodomain (EtM2e)

To generate a VSVΔG vector coexpressing the codon-optimized genes encoding EΔM-tM2e, H5-2005 (H5_05_), or H5-2024 (H5_22_), each of these cDNAs was cloned and inserted into an rVSVΔG vector, as indicated in Fig. 1A, and named VSV-EtM2e/H5_05_ or VSV-EtM2e/H5_22_. EΔM-tM2e (EtM2e) was generated by using Ebola glycoprotein DC-targeting domain (EΔM) fusion protein technology ^25^ through the insertion of four copies of the influenza M2 ectodomain (24 aa) polypeptide (tM2e, composed of two copies from human flu strains, one from a swine flu strain and one from an avian flu strain) (Fig. 1B, top panel) into EΔM to increase the immunogenicity of tM2e ^16^. In both the H5_05_ and H5_22_ open reading frames, mutations were introduced at the furin-dependent cleavage site to convert the furin-dependent cleavage site to a trypsin-dependent activation site characteristic of low-pathogenicity influenza envelope glycoprotein ^26^ (Fig. 1B, lower panel).

After these VSV vectors were constructed, each vector was rescued to obtain the vaccine candidates, V-EtM2e/H5_05_ or V-EtM2e/H5_22_, in Vero E6 cells via reverse genetics technology ^27^. Each vaccine candidate was subsequently tested for infection in Vero E6 cells. After 28 hours of infection, the infected cells were collected to detect different immunogens in the cell lysates. High expression of H5_05_, H5_22_, and EtM2e in each corresponding type of VSV-infected cell was detected by western blotting (WB) using antibodies specific for HA and M2 (Fig. 1C). The expression of the VSV nucleocapsid (N) protein was also detected in the VSV-infected cells (Fig. 1C, lower panel, lanes 1-2). Immunofluorescence assays confirmed the presence of both the HA and M2 proteins in the infected cells (Fig. 1D). These results revealed the concomitant expression of two avian influenza membrane proteins, HA and M2, in cells infected with the VSV vector vaccine candidates.

### 2. V-EtM2e/H5_05_ or V-EtM2e/H5_22_ induced robust anti-HA and anti-M2e IgG antibody responses in immunized mouse serum

In this study, the humoral immune responses induced by V-EtM2e/H5_05_ and V-EtM2e/H5_22_ were investigated by intramuscularly immunizing BALB/c mice with 1×10^7^ TCID_50_ of each vaccine product following a boost on day 17 (Fig. 2A). Sera from immunized mice were collected on days 14 and 28 postimmunization, and the levels of IgG antibodies specific to the HA of A/Cambodia/NPH230032/2023 (H5N1) or M2e (human) were determined by ELISA. On day 14, V-EtM2e/H5_22_**-**vaccinated mice had high anti-HA (1:300) and anti-M2 (1:500) IgG antibody endpoint titers (Fig. 2B). On day 28 (11 days after boost), the serum anti-HA and anti-M2 IgG antibody endpoint titers in the mouse serum pool were significantly greater (anti-HA, 1:3700; anti-M2, 1:14754) (Fig. 2B, right panel). We also evaluated the specific antibody levels in the serum from individual V-EtM2e/H5_22_- or V-EtM2e/H5_05_-vaccinated mice (28 days), and high levels of anti-HA and anti-M2 antibodies were detected in each immunized mouse (Fig. 2C). Anti-HA IgG subtypes in mouse serum were also quantified by analyzing T helper type 1 (Th1)-dependent antibody (IgG2a) and T helper type 2 (Th2)-dependent antibody (IgG1) levels. Interestingly, antibody levels, both V-EtM2e/H5_22_ and V-EtM2e/H5_05_ induced significantly higher levels of anti-HA IgG2a antibodies than anti-HA IgG1 antibodies (Fig. 2D). The above results indicate that these vaccine candidates are immunogenic and can induce efficient anti-HA and anti-M2e humoral immune responses.

**Figure 2.**
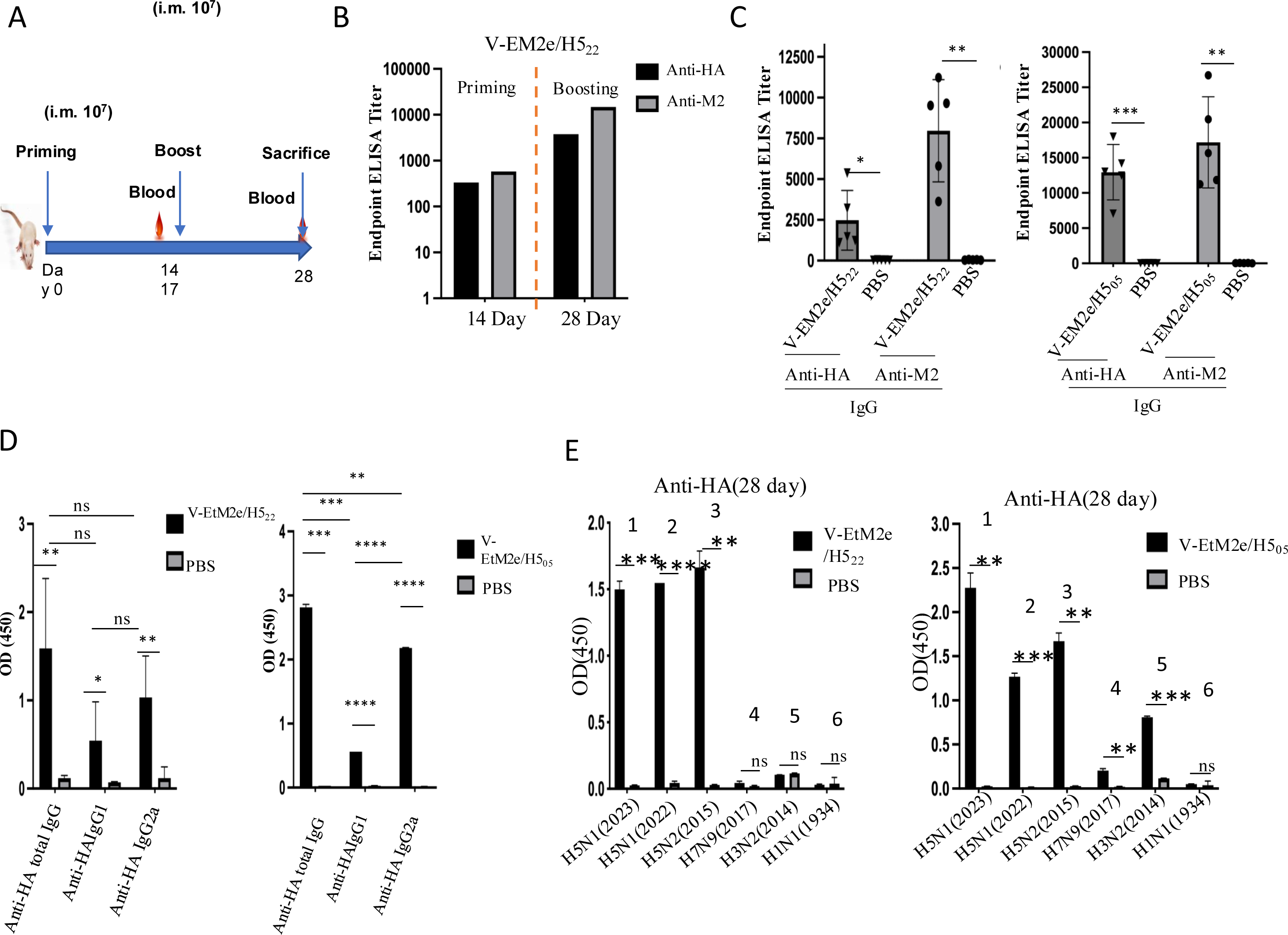
Anti-influenza HA and anti-M2e immune responses induced by rVSV vaccine candidates. A) Schematic of the immunization of V-EtM2e/H5_05_ or V-EtM2e/H5_22_ via the intramuscular (IM) route in BALB/c mice. B) The V-EtM2e/H5_22_ immunized mice sera were collected on days 14 and 28 post-immunization and measured for anti-HA and anti-M2 total IgG by ELISA. C) The endpoint titers of anti-HA and anti-M2 total IgG in V-EtM2e/H5_22_ (left panel) or V-EtM2e/H5_05_ (right panel) immunized individual mouse serum. D) The anti-HA IgG, IgG1 and IgG2a antibody levels in V-EtM2e/H5_22_ (left panel) or V-EtM2e/H5_05_ (right panel) immunized mice sera (1:200 dilution). E) The 28th day mouse sera pools of V-EtM2e/H5_05_ (right panel) or V-EtM2e/H5_22_ (left panel) were tested to determine if they can specifically recognize the recombinant HA proteins derived from different subtypes of influenza viruses, including H5N1, H5N2, H7N9, H3N2 and H1N1 by ELISA. Data represent mean ±SD. Statistical significance was determined using an unpaired T-test. *, p <0.05; **, p <0.01; *** p <0.001; ****, p <0.0001.

### 3. Specific recognition of different HA and M2 proteins from various influenza subtypes by vaccine-immunized sera

Next, we analyzed whether V-EtM2e/H5_22_ and V-EtM2e/H5_05_ vaccine-induced anti-HA antibodies could recognize recombinant HAs from different influenza subtypes, including H5 (recombinant HA synthesized through H5 genes isolated in different years), H1, H3, and H7 subtypes. The ELISA results revealed that both V-EtM2e/H5_22_ (Fig. 2E, left panel) and V-EtM2e/H5_05_ (Fig. 2E, right panel) immunization induced high levels of antibodies recognizing HA proteins from H5N1 (2022 and 2023) and H5N2 (2015) (Fig. 2E, bars 1-3). However, both types of vaccine-immunized sera, especially V-EtM2e/H5_22_-immunized sera, exhibited weak cross-reactive responses to H3 (2014) and/or H7 (2017) (Fig. 2E, bars 4 and 5) and very low binding to H1 (1934) (Fig. 2E, bar 6). Taken together, V-EtM2e/H5_05_ or V-EtM2e/H5_22_ immunization in mice induced a robust humoral response to HA from several H5N1 strains, as described here, but did not effectively recognize the HAs from the H1, H3 and H7 subtypes tested in this study.

### 4. Mouse anti-HA sera from V-EtM2e/H5_05_- or V-EtM2e/H5_22_-Immunized mice induced high levels of specific neutralizing antibodies against H5-pseudovirus infection

To evaluate whether immunization with V-EtM2e/H5_05_ or V-EtM2e/H5_22_ can induce protective immune responses against avian influenza H5-mediated viral infection, we tested neutralizing antibody (NAb) levels by using the H5-pseudotyped lentiviral virus (H5-PVLP) system in 293T cells, as described in the Materials and Methods. First, we analyzed whether each vaccine-immunized mouse serum sample could produce neutralizing antibodies against the corresponding H5 pseudovirus infection. The results clearly showed that all the V-EtM2e/H5_22_- and V-EtM2e/H5_05_-immunized mouse serum samples exhibited high neutralizing activity against the corresponding H5-PVLP infection (Fig. 3A). Conversely, the sera from V-EtM2e/H5_05_-immunized mice could not efficiently block H5_22_-PVLP infection, while the sera from V-EtM2e/H5_22_-immunized mice failed to effectively block H5_05_-PVLP infection (Fig. 4B). These findings substantially confirmed the strain specificity of NAbs induced by HA. Furthermore, we tested whether anti-M2e could enhance the anti-HA neutralization of M2^+^/H5_22_-PVLPs by testing the neutralization antibody ID50 titers in V-EtM2e/H5_22_-immunized mouse serum against M2^+^ or M2^-^/H5_22_-PVLPs. All individual mouse sera inhibited M2^+^/H5_22_-PVLP infection and M2^-^/H5_22_-PVLP infection at similar levels (Fig. 3C), suggesting that the presence of anti-M2e antibodies in the serum did not affect the neutralizing activity mediated by anti-HA antibodies.

**Figure 3.**
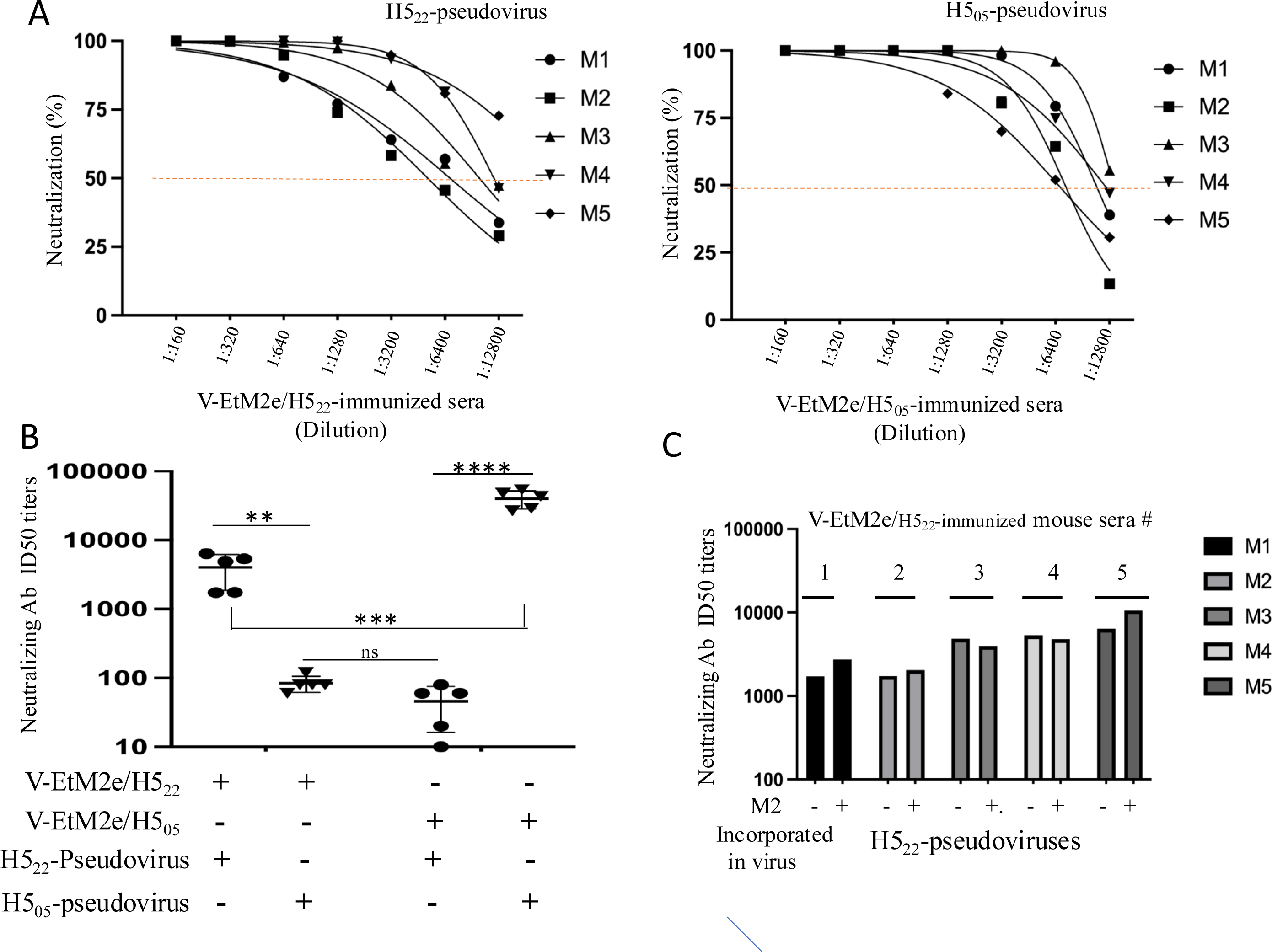
V-EtM2e/H5_05_ and V-EtM2e/H5_22_ vaccine candidates elicited neutralization antibodies. A) Representative neutralization curves in individual immunized mouse serum against H5_22_- or H5_05_-Luc-pseudovirus. Normalized percentage neutralization values from V-EtM2e/H5_22_ (left) or V-EtM2e/H5_05_ (right) vaccinated mouse sera and plotted against the dilution factors. M1-M5: each of the individual immunized mice. B) The ID50 titer of neutralizing antibody in each immunized mouse serum against H5_05_-Luc-pseudovirus or H5_22_-Luc-pseudovirus. C) The ID50 titer of neutralizing antibody in V-EtM2e/H5_22_ vaccine-immunized mouse serum pools against H5_22_-Luc-pseudovirus incorporated with or without M2 protein (as indicated). Data represent mean ±SD and were obtained from over two independent experiments. Statistical significance was determined using an ordinary one-way ANOVA and Turkey’s test. *, p <0.05; **, p <0.01; *** p <0.001; ****, p <0.0001.

**Figure 4.**
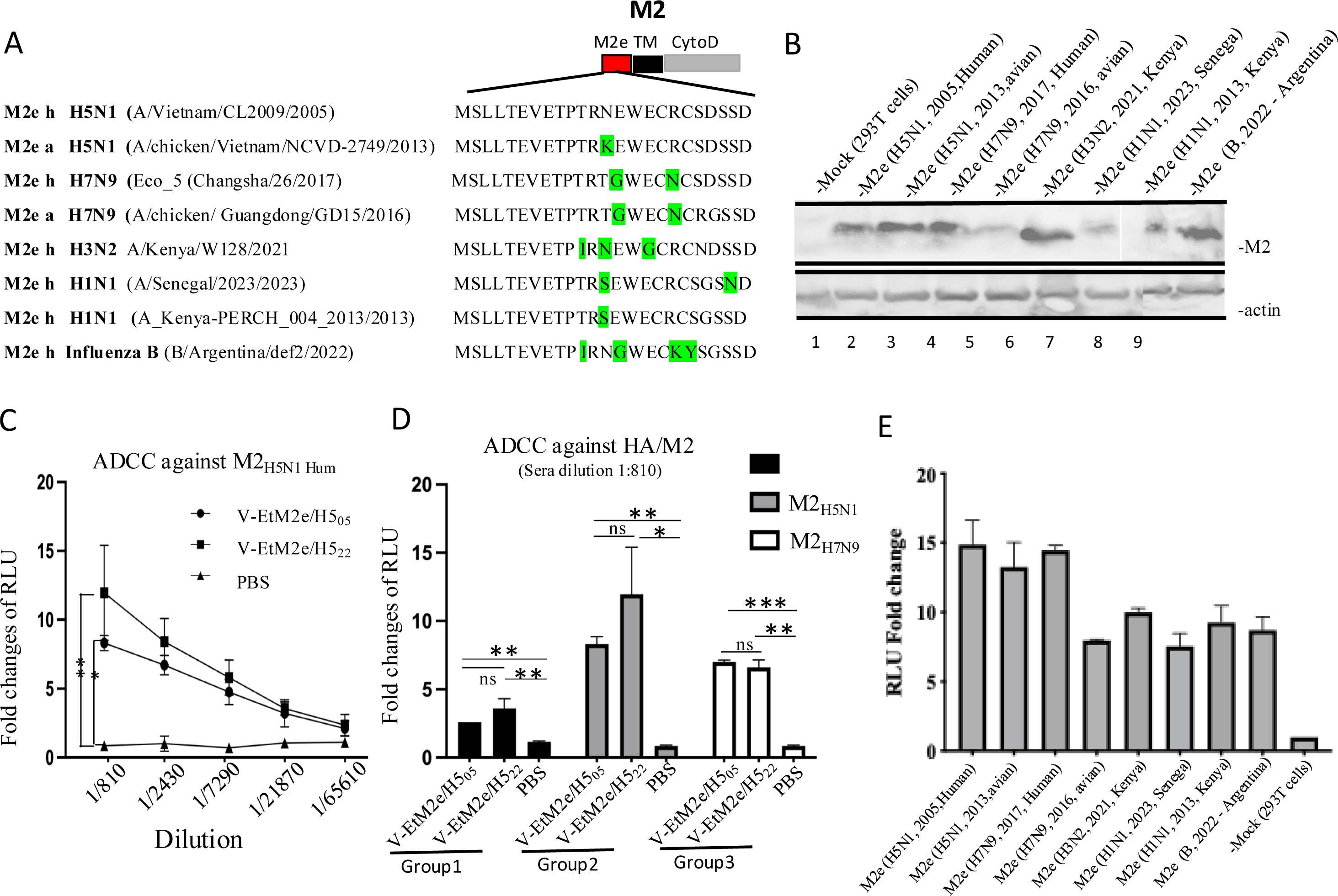
Vaccine immunization induces the ADCC activities against HA and M2e from a variety of influenza A subtypes. A) Sequence alignments of the ectodomain of M2 proteins (M2e) from different subtypes of influenza A and B. Each M2e encoded gene sequence was fused in frame to the transmembrane domain and the cytoplasmic region of an H5N1 M2 expressing plasmid (on the top of right panel). The variable amino acids in M2e, which are different from H5N1 (human, 2005), are marked in green. B) Each M2 plasmid encodes M2e from different subtypes of influenza (as indicated) was transfected into 293T cells. After 48 hours, the cells were lysed and run on a 12% SDS-PAGE gel, followed by immunoblotting with a pool of sera from V-EtM2e/H5_22_ immunized mice and an anti-actin antibody. C) The anti-HA ADCC activity induced by V-EtM2e/H5_05_ or V-EtM2e/H5_22_ immunized mice pooled sera (3×serially dilution) was detected by measuring the Luciferase activity from activated effector Jurkat cells and calculated as fold change against the no-serum control. D) Comparison of the anti-H5_22_ (group1), -M2e_H5_ (group2), and - M2e_H7_ (group 3)-ADCC activity levels mediated by pooled sera (1:810 dilution) from V-EtM2e/H5_05_- or V-EtM2e/H5_22_-immunized mice. E) V-EtM2e/H5_22_-immunized mice (1:810 dilution) mediated broad ADCC activities against each M2e from variable subtypes of influenza A and B, which was expressed on the transfected target 293T cells (as indicated).

### 5. Vaccination induced broad ADCC activity against different influenza M2 proteins

ADCC against influenza A virus infection has been shown to provide cross-strain protection and significantly contributes to viral clearance and the termination of viral infection ^28,29^. We therefore evaluated whether V-EtM2e/H5_05_ or V-EtM2e/H5_22_ vaccination could induce ADCC activity against different subtypes of influenza M2e by using an ADCC reporter system, as described in the Materials and Methods ^30,31^. To do so, we first constructed M2 plasmids expressing the gene encoding the M2 ectodomain from different strains, including H5N1 (human and avian), H7N9 (human and avian), H3N2 (2021), H1N1 (2023 and 2013), and influenza B (2022 and Argentina) ^32^, which were fused with the transmembrane domain and the cytoplasmic region of H5N1 ^21^ (Fig. 4A). To confirm the expression of different subtypes of M2e, each M2 expressor was transfected into 293T cells, after which the cells were lysed and processed for western blot analyses by using V-EtM2e/H5_22_-induced anti-M2e antibodies in the serum of vaccinated mice. The results revealed that the serum of vaccinated mice can recognize the M2 ectodomain from a wide range of influenza A strains with variable binding affinities (Fig. 4B). Such broad binding ability of the anti-M2e antibody from the vaccinated mice is expected since the ectodomain of M2 is highly conserved among viruses, and our V-EtM2e/H5_22_ vaccine contains four copies of the M2 ectodomain polypeptide, including two copies from human flu strains, one from a swine flu strain and one from an avian flu strain (Fig. 1A and B).

We next examined the ADCC activity mediated by immunized mouse serum against each influenza subtype M2e. Dose-dependent analyses revealed that both V-EtM2e/H5_05_- and V-EtM2e/H5_22_-immunized mouse serum pools induced strong ADCC activity against M2 from H5_05_ on the surface of transfected 293T cells in a dose-dependent manner (Fig. 4C). Additionally, we compared the anti-HA and anti-M2e ADCC response levels of two immunized mouse serum samples, and the results revealed that compared with the no serum control, both the V-EtM2e/H5_05_-and the V-EtM2e/H5_22_-immunized mouse serum pools induced ADCC activity against H5_22_, with 2–4-fold increases at a dilution of 1:810 (fold change of 0) (Fig. 4D, group 1). Interestingly, both immunized mouse serum pools induced significantly greater ADCC activity against M2_H5N1_- and M2_H7N9_-expressing cells (8- to 14-fold and 6- to 7-fold, respectively) (Fig. 4D, groups 2 and 3). Furthermore, we investigated the V-EtM2e/H5_22_-induced ADCC against M2e from different influenza viruses. The results revealed that M2 antibodies in mouse serum can mediate similarly high levels of ADCC activity against distinct influenza subtypes (Fig. 4E). Taken together, these results provide evidence that vaccination with either V-EtM2e/H5_05_ or V-EtM2e/H5_22_ can elicit remarkable levels of anti-HA and M2 antibodies that can mediate ADCC activity against different subtypes of influenza viruses.

### 6. Immunization with V-EtM2e/H5_22_ via either the IN or IM route efficiently protects mice from lethal H5N1 influenza virus infection

To investigate the protective potential of the V-EtM2e/H5_22_ vaccine against H5N1 influenza virus infection, BALB/c mice (*n* = 40, 20 females and 20 males) were immunized with V-EtM2e/H5_22_ via the IM (1 × 10^7^ TCID_50_) or IN (1 × 10^5^ TCID_50_) route. As a control, VSV-EBOVGP was administered to mice (n = 40) at the same dose and via the same route. Two weeks after the boost vaccination, the mice were intranasally challenged with a 10× lethal dose (50% (LD_50_)) of the H5N1 virus (A/BC/PHL_2032/2024). Half of the mice (5 mice/group) were euthanized and necropsied at 4 days post-infection (dpi), at which time serum, lung, heart, kidney and spleen samples were collected to quantify the infectious H5N1 viral titer in the tissue homogenates (Fig. 6A). The remaining animals were weighed and monitored daily until 18 dpi. We found that both IM and IN administration of V-EtM2e/H5_22_ provided 100% protection against lethal H5N1 infection, whereas all the mice in the control groups died within 5 days (Fig. 6B and C). Very intriguingly, no viral burden was detected in the heart, kidney, spleen or lungs of IM- and IN-vaccinated mice after challenge with H5N1. In mice immunized with the VSV-EBOVGP vaccine, H5N1 infection resulted in high H5N1 viral loads in different tissues, the highest levels of which were detected in the lungs (Fig. 6D). These results indicated that immunization with V-EtM2e/H5_22_ via either the IN or the IM route completely protected mice from lethal H5N1 influenza virus challenge.

## Discussion

A multivalent influenza vaccine capable of protecting against diverse influenza A strains, particularly highly pathogenic avian influenza (HPAI) strains, has been a central objective of influenza research for decades. In this study, we developed recombinant VSV-based vaccine candidates that coexpress HPAI H5 hemagglutinin (HA) and four copies of the influenza matrix protein 2 ectodomain (M2e), which were fused to the dendritic cell-targeting domain of Ebola glycoprotein (EΔM). Our findings illustrate that these candidates elicited strong immune responses against both HA and M2e, resulting in high neutralizing activity and robust antibody-dependent cellular cytotoxicity (ADCC) against different influenza subtypes. Most importantly, the immunized mice were fully protected against lethal challenge with HPAI H5N1, highlighting the protective potential of this multivalent vaccine strategy.

Our previous studies demonstrated that the insertion of large heterologous polypeptides into the mucin-like domain (MLD) of the Ebola glycoprotein (EΔM) allows for the efficient delivery of antigens to human macrophages and dendritic cells to induce robust immune responses ^25,33^. Building on this strategy, we incorporated four copies of M2e from different influenza strains (two from human strains, one from an avian strain, and one from a swine strain) into the EΔM scaffold to generate an EΔM-tM2e construct, as previously described ^16^ (Fig. 1A), which was designed to increase immune recognition of this conserved M2e antigen. In parallel, full-length cleavage-mutated H5_05_ or H5_22_ HA immunogens were inserted into the VSV vector, producing two vaccine candidates (V-EtM2e/H5_05_ and V-EtM2e/H5_22_) that simultaneously expressed both HA and M2e. Western blot and fluorescence analyses confirmed the efficient expression of H5_05_, H5_22_, and M2e in Vero-E6 cells infected with either construct (Fig. 1B and C). Consistently, animal immunization studies clearly showed that these vaccines induced robust anti-HA and anti-M2e antibody levels (Fig. 2), demonstrating the high immunogenicity of the influenza antigens presented in this VSV-based vaccine platform.

We next examined the breadth of anti-HA responses induced by vaccination. Mouse sera strongly recognized recombinant H5 proteins from the H5N1 strains isolated from 2015, 2022, and 2023 but bound weakly to H3 (2014) and H7 (2017) and could not bind H1 (1934) well (Fig. 3). These findings suggest that the HA-directed immune response is largely subtype restricted. Given that HA mediates viral attachment and entry, neutralizing antibodies play a central role in blocking early infection. By using a luciferase-expressing H5-PVLP infection assay, sera from V-EtM2e/H5_22_-immunized mice exhibited strong neutralization of H5-PVLP infection but limited activity against H5-PVLPs, whereas the sera from V-EtM2e/H5_05_-immunized mice showed the opposite pattern (Fig. 4). Comparison of the H5_05_ and H5_22_ sequences revealed numerous mutations, particularly within the HA head domain, which governs receptor binding (Fig. S1). These differences likely explain the limited cross-neutralization. Such findings underscore the importance of accounting for variability in the HA head when HA-based vaccines are designed to optimize neutralizing capacity. Several vaccine strategies seek to redirect immune responses toward the conserved HA stalk, thereby improving the breadth of the vaccine and reducing the need for annual reformulation ^34,35^. Our results suggest that enhancing stalk-directed immunogenicity with full-length HA vaccines remains important. Finally, although neutralizing antibody titers against heterologous H5 were relatively low, partial inhibition at higher serum concentrations was observed (Fig. 4). Whether such low neutralization is sufficient to confer in vivo protection warrants further investigation.

Another vital component of our vaccine design is the highly conserved ectodomain of the influenza matrix protein 2 (M2e), which is a promising universal influenza vaccine target because of its stability and limited sequence variation. Previous studies have shown that the ectodomain of the influenza virus M2 protein can mediate either antibody-dependent cellular cytotoxicity (ADCC) or antibody-dependent cellular phagocytosis (ADCP), which leads to the elimination of the influenza virus or the destruction of cells that are already infected ^36,37^. In this study, in addition to the H5 immunogen in the vaccine, four copies of influenza M2e (two from human strains and one each from avian and swine strains) were expressed in our vaccine. To increase the immunogenicity of M2e, tM2e was fused with EΔM, which can efficiently target DCs and macrophages, resulting in significant increases in the production of M2e-specific immune responses ^16^. Indeed, the anti-M2e immune responses induced by V-EtM2e/H5_05_ and V-EtM2e/H5_22_ were not only able to recognize M2e derived from H5N1 but also bind to M2e derived from human and avian H7N9 strains and to H1N1, H3N2 and influenza B (Fig. 3C). Notably, vaccine-immunized sera were able to mediate strong ADCC activity against M2e derived from H5N1, H7N9, H3N2 and H1N1 subtypes (Fig. 5D), suggesting potential broad protection of these vaccines against different influenza A infections from various host ranges (human, avian and swine). Our previous study demonstrated that immunization with EΔM-tM2e via the intramuscular or intranasal route provides 100% protection against lethal H1N1 and/or H3N2 influenza virus challenge ^16,22,38^. Further investigations are underway to test whether V-EtM2e/H5_22_ could mediate broad protection against infections by other human and avian influenza A viruses.

**Figure 5.**
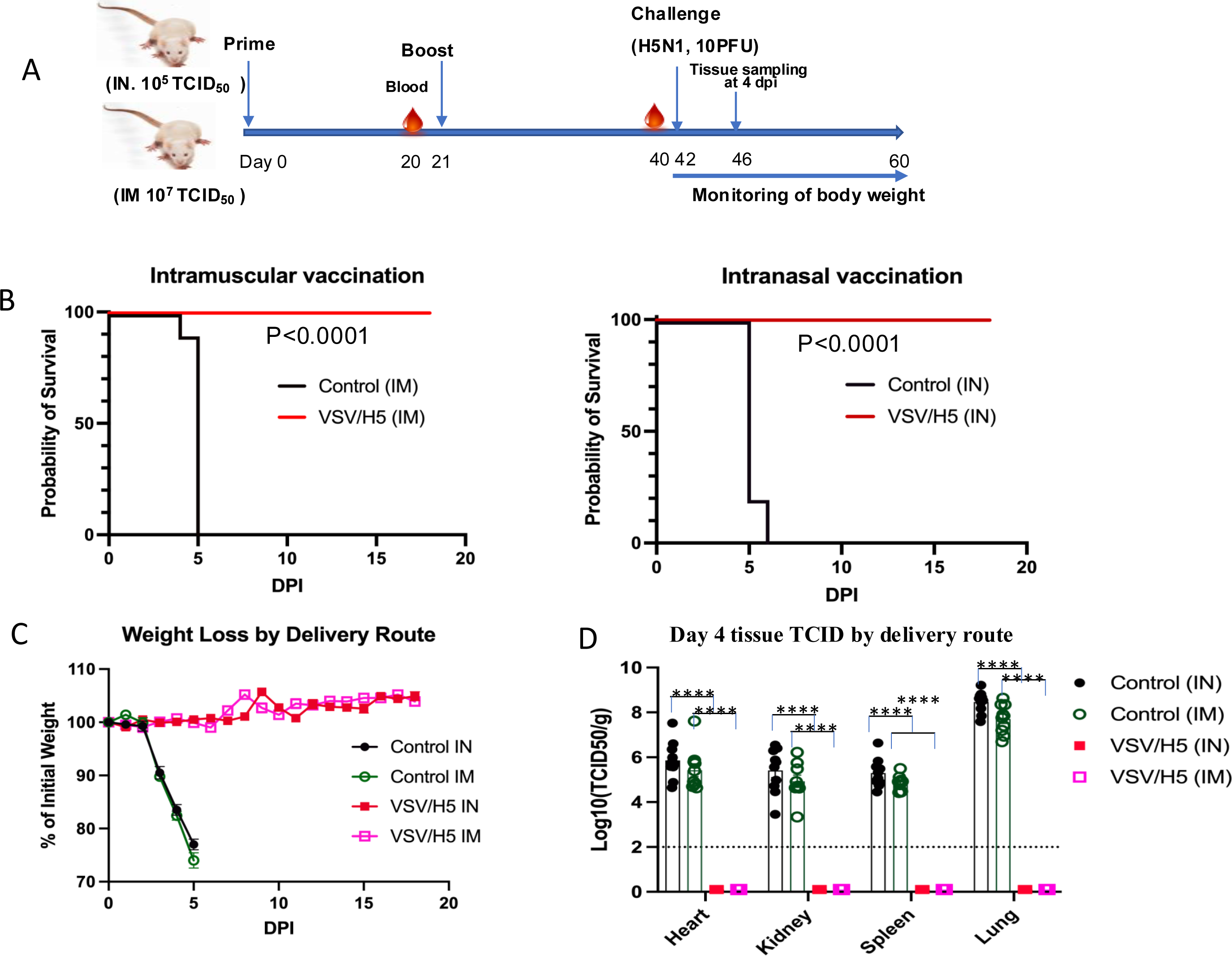
V-EtM2e/H5_22_ protected against lethal H5N1 influenza virus infection in mice. A) Schematic of the VSV vaccine candidate immunization and H5N1 influenza challenge protocol used in the study. Briefly, BALB/c mice were administered with V-EtM2e/H5_22_ or VSV-EBOVGP (control) via IM or IN at day 0 and day 21. Two weeks after the boost vaccination, the vaccinated mice were intranasally challenged with H5N1 virus (A/BC/PHL_2032/2024). Animals were weighed and monitored daily, half of the mice were euthanized and necropsied 4 days post-infection, and their serum, lung, heart, kidney and spleen samples were collected for quantifying the infectious H5N1 virus in tissue homogenates. B, C) Survival and weight loss in the vaccinated or the control mice following infection with the H5N1 virus. D) Viral burden in various tissues was detected by measuring the TCID_50_ values per gram of tissue in MDCK cells that were infected by tissue homogenates from challenged mice. n =20 for B and C, 20 through day 40, and 20 at day 42; n =10 for D (from day 4 post-infection). Statistical significance was assessed by two-way ANOVA analysis with multiple comparisons (A, B) and the Mantel-Cox test (D). *, p <0.05; **, p <0.01; *** p <0.001; ****, p <0.0001.

We evaluated the protective efficacy of V-EtM2e/H5_22_ against lethal H5N1 challenge in mice. Both intramuscular (IM) and intranasal (IN) immunization conferred complete protection to male and female animals following exposure to a highly pathogenic H5N1 strain isolated in 2024 (Fig. 6). None of the immunized mice exhibited significant weight loss after infection, in stark contrast to the control animals. Notably, at four days post-challenge, the infectious H5N1 virus was not detected in multiple tissues, including the lungs, spleen, heart, and kidney, of the immunized mice compared with tissues from the control groups (Fig. 6D). These findings provide compelling evidence that V-EtM2e/H5_22_ (delivered via either the IM or IN route) can effectively prevent currently circulating H5N1 infection, clinical disease, and systemic viral replication in a mouse model. Together, the results of this study strongly support the further investigation of this vaccine platform for broader translational applications.

Since the mucosal membranes of the respiratory tract serve as the natural entry point for influenza viruses, these membranes represent the first line of host defense. Ideally, compared with intramuscular vaccination, mucosal vaccination via the respiratory tract may elicit stronger local protective immune responses. The results of this study demonstrated that IN administration, even at 100-fold lower doses, achieved protection equal to that of IM administration (Fig. 6). This further supports the high efficacy of noninvasive mucosal delivery with significantly lower doses of the vaccine to block influenza infection. It should be noted that, the trypsin-dependent sensitive site of H5 in our vaccine platform resulted in the expression of a low-pathogenic influenza envelope glycoprotein (Fig. 1B) ^26^, which can improve the safety profile without impairing the immunogenicity of the vaccine candidate. Together, these studies provide evidence for the high efficacy and attenuated feature of this bivalent vaccine platform, which can be easily adapted to generate effective influenza vaccines against different strains of influenza virus.

Overall, the results of this study demonstrate that the V-EtM2e/H5_05_ and V-EtM2e/H5_22_ vaccines elicited strong influenza-specific immune responses and provided complete protection against lethal HPAI H5N1 challenge. By integrating HA, a potent but antigenically variable target, with M2e, a highly conserved epitope that mediates Fc-dependent effector functions, this platform achieves both subtype-specific neutralization and broader cross-strain protection. These findings highlight the promise of rVSV-based multivalent vaccines as a foundation for developing a universal influenza A vaccine. Future studies will be essential to evaluate the durability of protection, assess efficacy against additional influenza strains, and validate performance in nonhuman primate models.

## Data Availability Statement

The data used in this study are available in the main text. The corresponding author can be contacted for additional information.

## Ethics Statement

The animal study was reviewed and approved by the University of Manitoba Animal Care and Use Committee.

## Author contributions

The experiments were conceived of and designed by Z.J.A., K.F., D.K., and X.J.Y. The constructions and rescue of different rVSV vaccine candidates and the characterization of viral replication were performed by Z.J.A. and X.J.Y. Animal immunization studies were conducted by Z.J.A. S.A., M.B., D.L., N.G., P.L., and M.N. The Influenza H5N1 challenge experiments and related analyses were carried out by R.V., T.T., and D.K. The initial draft of the paper was written by Z.J.A., N.G., R.V., and X.J.Y. All other authors contributed to editing the paper into its final form. The work was managed and supervised by D.K., and X.J.Y.

## Acknowledgements

We thank Drs. David C. Alexander, Shawn Babiuk and Anthony Signore for their technical support. M. Buyu is the recipient of the Canadian International Development Scholarship (BCDI2030) funded by Global Affairs Canada (GAC). This research was supported by the Operating Grants from the Canadian Institutes of Health Research (CIHR) (OS1-190775) and Social Sciences and Humanities Research Council of Canada (SSHRC) (CBRF2-2023-00217).

---

**Fig. S1.**
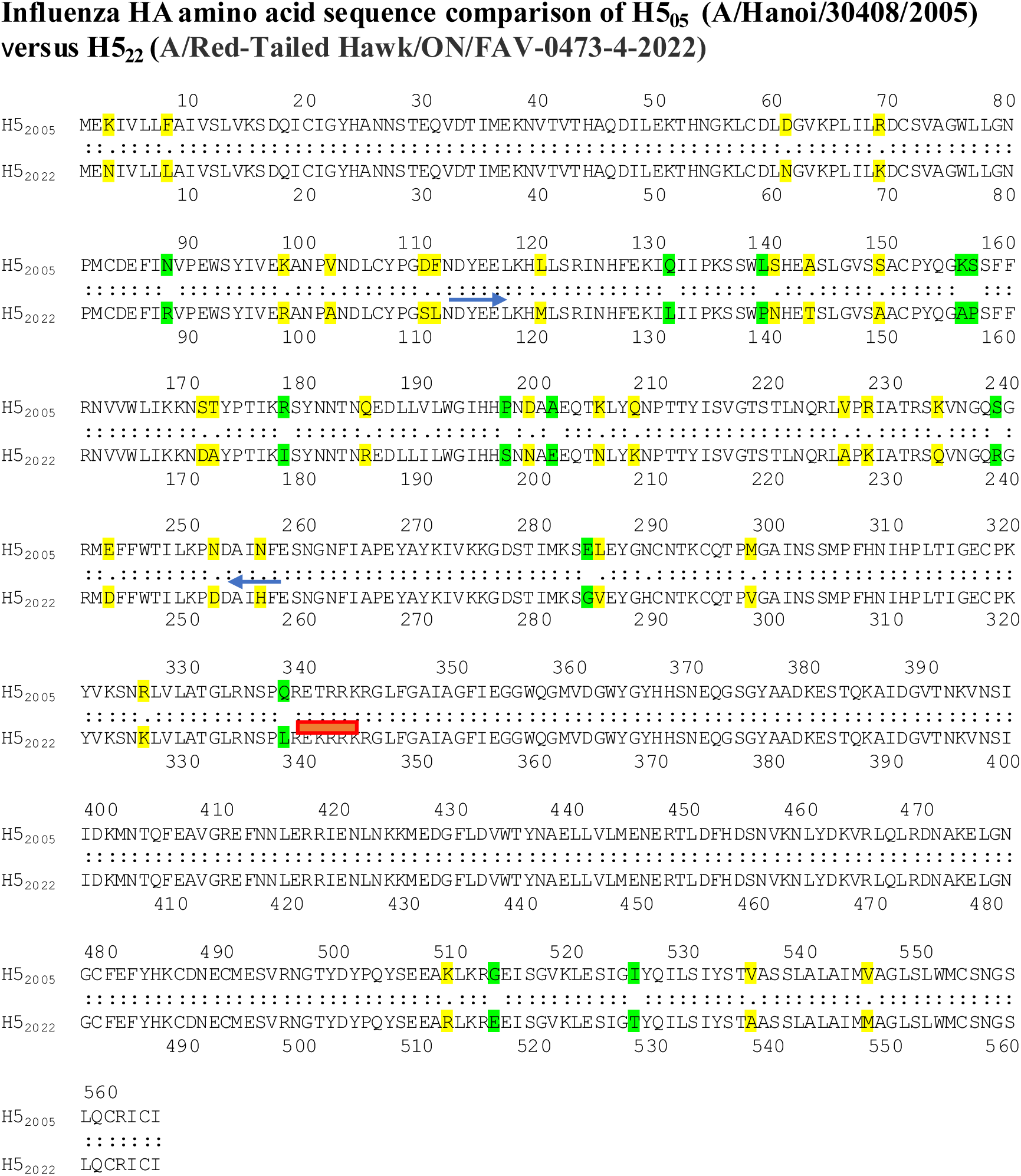
Amino acid (aa) alignment of HA05 and HA22. The blue arrows show the head region, and the red line indicates the cleavage site. The different aa are marked in green and yellow

